# Tailer: A Pipeline for Sequencing-Based Analysis of Non-Polyadenylated RNA 3’ End Processing

**DOI:** 10.1101/2021.12.06.471174

**Authors:** Tim Nicholson-Shaw, Jens Lykke-Andersen

## Abstract

Post-transcriptional trimming and tailing of RNA 3’ ends play key roles in the processing and quality control of non-coding RNAs (ncRNAs). However, bioinformatic tools to examine changes in the RNA 3’ “tailome” are sparse and not standardized. Here we present Tailer, a bioinformatic pipeline in two parts that allows for robust quantification and analysis of tail information from next generation sequencing experiments that preserve RNA 3’ end information. The first part of Tailer, Tailer-Processing, uses genome annotation or reference FASTA gene sequences to quantify RNA 3’ ends from SAM-formatted alignment files or FASTQ sequence read files produced from sequencing experiments. The second part, Tailer-Analysis, uses the output of Tailer-Processing to identify statistically significant RNA targets of trimming and tailing and create graphs for data exploration. We apply Tailer to RNA 3’ end sequencing experiments from three published studies and find that it accurately and reproducibly recapitulates key findings. Thus, Tailer should be a useful and easily accessible tool to globally investigate tailing dynamics of non-polyadenylated RNAs and conditions that perturb them.

## Introduction

Dynamic post-transcriptional addition and removal of nucleotides from the 3’ ends of RNAs is a key hub for RNA maturation and regulation. While these dynamics are perhaps best understood for eukaryotic mRNAs that undergo polyadenylation (Lee et al. 1971; Darnell et al. 1971; Edmonds et al. 1971) and deadenylation (Goldstrohm and Wickens 2008) to regulate translation and stability (Nicholson and Pasquinelli 2019), non-coding RNAs also experience a wide variety of 3’ end modifications. These events, which include 3’ end trimming, tailing, or chemical modification (Yu and Kim 2020; Liudkovska and Dziembowski 2021; Perumal and Reddy 2002), can have different functional consequences depending on the RNA, the modification, and the cellular context. Some 3’ end modification events, exemplified by CCA addition to tRNAs (Deutscher 1973), play key roles in RNA maturation and serve to produce mature 3’ ends that promote RNA stability and/or function (Dupasquier et al. 2008; Katoh et al. 2009; Nguyen et al. 2015; Shukla and Parker 2017). Other modifications promote rapid degradation, for example as part of quality control pathways that detect and degrade aberrant or damaged transcripts (Shcherbik et al. 2010; Liu et al. 2014; Lardelli and Lykke-Andersen 2020; LaCava et al. 2005). These processes are essential to life and their dysfunction can lead to human disease (Wolin and Maquat 2019); yet, how enzymes acting at RNA 3’ ends cooperate and compete to dictate RNA function and stability remains poorly defined for the majority of RNAs.

Early characterizations of non-coding (nc)RNAs and their 3’ end sequences focused on single RNA species, initially using radioisotope labeling and enzymatic digestions, and later, RNA 3’ end amplification methods coupled with cloning and sequencing (Lund and Dahlberg 1992; Rinke and Steitz 1982; Frohman et al. 1988). More recent advances in sequencing technology have allowed for examination of ncRNA ends on a transcriptome-wide level, and for monitoring how those ends change globally in response to perturbations. Techniques such as ligation-based 3’ Rapid Amplification of cDNA Ends (3’RACE) coupled with high-throughput sequencing (Shukla and Parker 2017; Lee et al. 2014) can provide a snapshot at nucleotide level resolution of RNA 3’ ends globally. A typical reverse genetics approach to understanding RNA 3’ end dynamics involves identifying enzymes capable of modulating RNA tails, depleting them from cells, and monitoring changes in RNA 3’ ends, thereby identifying potential direct targets of those enzymes (Łabno et al. 2016; Lardelli et al. 2017; Son et al. 2018; Lardelli and Lykke-Andersen 2020; Allmang et al. 1999; Berndt et al. 2012). During data analysis, changes to RNA 3’ ends are generally quantified with scripts and pipelines individual to each lab. While many of these scripts are made publicly available (for example (Pirouz et al. 2019)), easy-to-use and generalizable tools have been missing to make these types of analyses accessible to the broader research community.

Here we present Tailer, an easy to use and open-source pipeline that can analyze the status and perturbations of non-coding RNA 3’ ends from sequencing datasets for which RNA 3’ ends have been preserved. Tailer is fully featured, easily installable, and allows for analysis of new and previously published datasets. This pipeline takes mapped SAM or BAM files from 3’ end sequencing experiments, globally identifies positions and compositions of RNA 3’ ends, including their post-transcriptional tails, and outputs the data into a human readable CSV format. This output CSV file can then be uploaded to a web server, which provides utilities to discover RNAs undergoing statistically significant changes at their 3’ ends and to visualize RNA tail dynamics. The pipeline also allows for analysis of individual RNAs of interest from global or gene-specific sequencing experiments using local alignment.

To validate Tailer, we reanalyzed publicly available global and gene-specific 3’ end sequencing datasets from three studies focused on the exonucleases DIS3L2, TOE1 and PARN in human cells (Łabno et al. 2016; Son et al. 2018; Lardelli and Lykke-Andersen 2020). In all cases, Tailer identified target RNAs highlighted in the studies and faithfully reproduced observed effects on RNA 3’ ends. This validates the utility of Tailer as a tool to monitor global and gene-specific 3’ end processing of non-coding RNAs. While applied here to human RNA sequencing datasets, the pipeline is compatible with datasets from any organism of interest with reliable annotation information.

## Results and Discussion

### Pipeline Overview

Tailer is comprised of two arms (Fig. 1), Tailer-Processing, which identifies and quantifies 3’ end compositions of non-polyadenylated RNAs from 3’ end sequencing data, and Tailer-Analysis, an R-based Shiny app for candidate discovery and data visualization. Tailer is written in Python 3 (Van Rossum G and Drake FL. 2019), can be installed using the Package Installer for Python (PIP) accessed from the PyPi index (detailed installation instructions can be found on the readme page), and can be run from the command-line. The output of Tailer-Processing is a comma separated values (CSV) file, hereafter referred to as a Tail CSV file, which lists the identity and quantity of all 3’ ends of RNAs observed in the analyzed 3’ sequencing experiment that match a given annotation file, or a given list of genes. The Tail CSV file can then be fed into the Shiny-based (Chang et al. 2021) Tailer-Analysis web application for further analysis.

**FIGURE 1.**
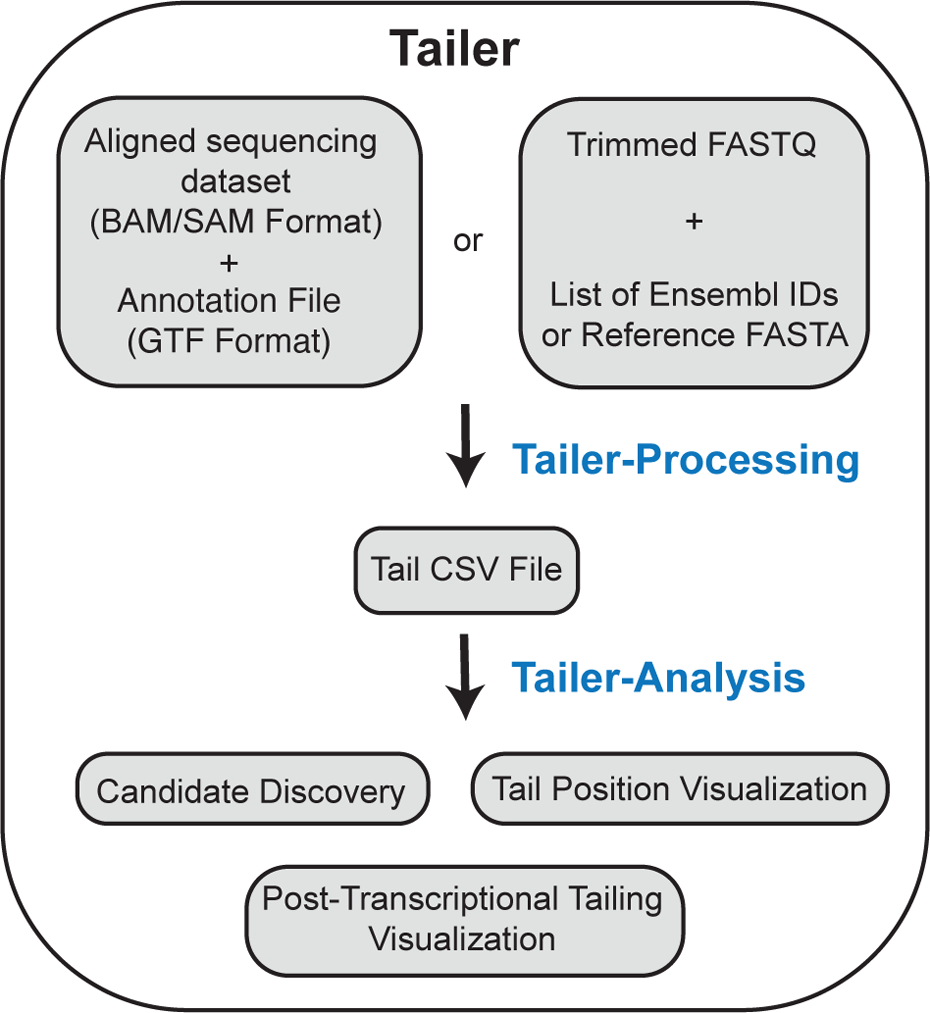
A general overview of Tailer’s workflow. Tailer is split into two major parts, a processing function, and an analysis webserver. Tailer-Processing infers RNA 3’ ends using a BAM/SAM alignment file and a GTF formatted annotation file, or a FASTQ sequence read file and a reference FASTA gene file (which can be generated from Ensembl IDs). For either method, the output is a standardized Tail CSV file, which can be analyzed directly, or fed into the Tailer-Analysis web server for discovery of candidate tailing changes in comparison between datasets as well as visualization of tails with a variety of graphing tools.

### Tailer-Processing in global mode annotates SAM/BAM files and calculates RNA 3’ end information

Tailer-Processing can be run from the command-line and can be used in a global mode to identify all RNA 3’ ends matching a genome annotation, or in local mode to identify 3’ ends of specific RNAs of interest (Fig. 2A). When used in global mode (Fig. 2B, left), Tailer-Processing requires two inputs, a SAM or BAM formatted alignment file and a GTF formatted annotation database. Experimentally, the sequencing data entered into Tailer-Processing should be generated using a library preparation method that preserves the 3’ end information of RNAs, such as a 3’ RACE (Frohman et al. 1988) experiment, and should be performed on RNA that is not poly(A)-selected. For small RNA 3’ end analyses, the RNA can be size selected prior to sequencing, but analyses can be performed on any sequencing experiment that preserves RNA 3’ ends.

**FIGURE 2.**
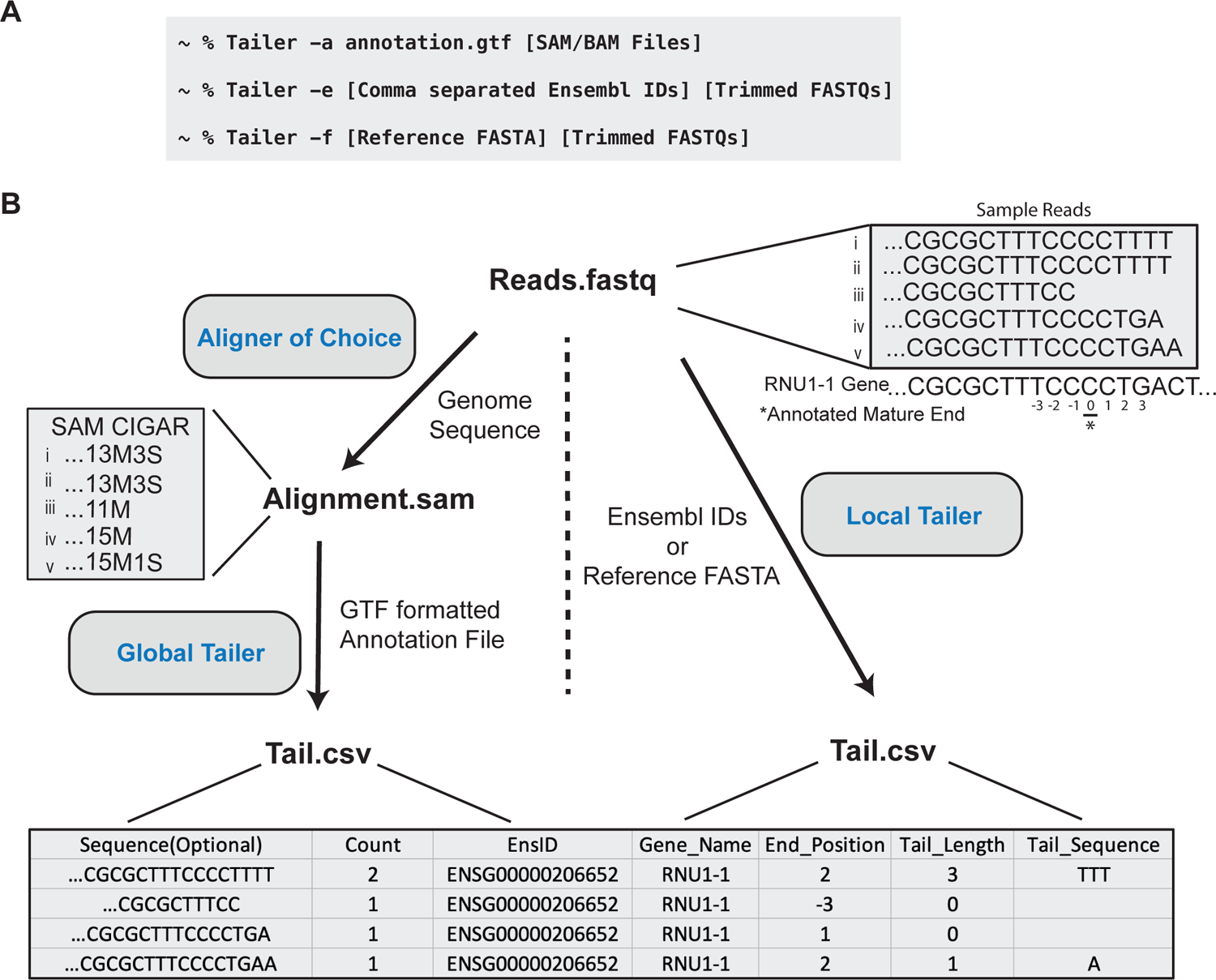
Tailer-Processing commands and examples of tail inference. (A) Example commands to run Tailer-Processing. After installation with PIP, Tailer-Processing can be invoked by typing “Tailer” into the command line. An -h flag will provide usage information that is also available in a readme.txt file. Examples of running Tailer-Processing in global mode, in local mode using Ensembl IDs, and local mode with a reference FASTA gene file are shown. (B) Global Tailer-Processing (left) uses SAM/BAM-formatted alignment files to infer RNA 3’ ends and post-transcriptional tails based on the SAM CIGAR and GTF annotation files. Local Tailer (right) does not require pre-alignment of the sequencing data and uses BLAST to align to a user provided FASTA gene database or one generated from provided Ensembl IDs. Local Tailer makes use of reported BLAST metrics of last query mapping position and last reference mapping position to infer tail information. Both modes produce a Tail CSV file with the same columns (bottom). An example output from five reads aligning to the RNU1-1 gene with corresponding SAM-file CIGAR strings is shown.

Sequencing can be performed either as single end reads from the 3’ end, or as paired end reads for improved alignment accuracy. Sequencing outputs, after trimming of any linkers, need to be subsequently aligned to a reference genome using any aligner that supports soft-clipping and produces a SAM/BAM-formatted alignment file output. It is important that the aligner supports soft-clipping as Tailer uses this feature to determine post-transcriptional tailing (for options that bypass soft-clipping, see below). Typically, we use STAR aligner (Dobin et al. 2013) with the following settings from Son, et al 2018 when interested in small non-coding RNAs (STAR --alignIntronMin 9999999 -- outFilterMultimapNmax 1000), which allows alignment to multicopy genes and disallows unannotated introns. The GTF file can be provided to Tailer as a full genome annotation or filtered to contain only genes of interest to create smaller sized output files. In case of paired-end sequencing, the specific read that corresponds to the RNA 3’ end needs to be specified with the “--read” flag. This pipeline has been most rigorously tested with annotations provided by the ENSEMBL database (Howe et al. 2021).

The output tail CSV file produced by Tailer-Processing reports the number of occurrences of each type of RNA 3’ end that is detected in the sequencing data (Fig. 2B, bottom). For each alignment reported in the input SAM alignment file, Tailer-Processing identifies the corresponding gene from the input annotation GTF file, identifies the 3’ end position of the read relative to the annotated gene 3’ end, and predicts any post-transcriptionally added tail. The gene, which is reported in the ‘Gene’ column, is identified as the gene in the same orientation as the aligned read that has the closest annotated 3’ end to the 3’-most aligned nucleotide of the read, with a requirement that the 3’ ends are within a default of 100 nucleotides of one another, an option that can be modified with the -t flag. To identify the read 3’ end position and predict any post-transcriptional tail, Tailer examines the CIGAR string reported in the SAM alignment file and searches for soft-clipping at the 3’ end of the read, i.e. the 3’ terminal nucleotides of the read that did not align to the genome (Fig. 2B, left). The composition of the soft-clipped nucleotides is reported as the post-transcriptional tail of the read in the output tail file (“Tail_Sequence” column). The position, including any post-transcriptional tail, of the last nucleotide of the read relative to the annotated 3’ end of the gene is reported as the 3’ end position of the read (“End_Position” column). For multi-mapped reads, all annotated genes aligning to the read are reported in the output file, which allows for more accurate downstream analyses of RNAs that are produced from multiple loci or have closely related pseudogenes (see below). In cases where reads align to multiple genes that are annotated with different 3’ ends, Tailer reports only the gene whose annotated 3’ end is closest to the calculated genome-encoded 3’ end of the read. Reads corresponding to identical RNA 3’ end sequences are finally collapsed and the number of reads for each is reported in the “Counts” column. This output format greatly reduces the size of the file to focus on information that is pertinent to tail analysis and can be reasonably uploaded to a web server. An optional column with 3’ end read sequences that can be useful for verification and troubleshooting purposes can be included in the tail file by implementing an -s flag.

### Running Tailer-Processing in local mode allows for rapid analysis of specific RNAs without the necessity for previous alignment or reliance on soft-clipping

Analysis by Tailer also lends itself to a gene-specific approach for greater depth on specific genes of interest using local alignments (Fig. 2B, right). This mode requires the user to have command line BLAST installed and a reference to it stored in the PATH variable on their workstation (https://www.ncbi.nlm.nih.gov/books/NBK279690/). The required inputs are a FASTQ file containing the called bases from the sequencing experiment, trimmed of any linkers, and one or more genes, either identified by their Ensembl IDs or provided in a FASTA file. For paired-end sequencing, the FASTQ file used should be the read file that corresponds to the 3’ end (typically read 2 for Illumina sequencing).

This mode can be used for analyses of gene-specific 3’ end sequencing data (Lardelli et al. 2017; Lardelli and Lykke-Andersen 2020). It can also be used for cases where soft-clipping is problematic for correct alignments (Suzuki et al. 2011), such as for genes that have closely related variants. In this case, initial global sequence alignments can be performed in the absence of soft-clipping and reads aligning to specific genes can subsequently be extracted from the SAM/BAM alignment files and converted back to FASTQ files with tools such as Bedtools (Quinlan and Hall 2010) and Samtools (Li et al. 2009). The local gene-specific Tailer can also be used directly on large FASTQ files from global sequencing experiments, but processing will be much slower than using global Tailer. The gene-specific mode downloads gene information from Ensembl along with 50 nucleotides of downstream sequence to aid in distinguishing between genome-encoded tails and post-transcriptionally added tails. Alternatively, a custom FASTA-formatted reference sequence can be provided instead with the -f flag (Fig. 2A). This reference sequence should contain genomic sequence downstream of the gene for accurate distinction between genomic-encoded and post-transcriptional tails. When including downstream sequence, the -m flag should be used to specify the number of downstream nucleotides included in the reference to ensure that the mature end is correctly annotated.

After building a BLAST formatted database with the downloaded sequences, gene-specific Tailer aligns each read in the FASTQ file and uses the alignment information to calculate the read 3’ end position relative to the annotated gene 3’ end and identify the composition of any predicted post-transcriptional tail, producing an output Tail CSV file identical to that produced by global Tailer-Processing. It is important to note that for both the global and local methods, post-transcriptional tails are predictions based on absence of alignment. These generally represent the most conservative predictions for the actual post-transcriptional tail, since any post-transcriptionally added nucleotide that matches a nucleotide encoded from the genome will be assigned as genome-encoded by default.

### Using Tailer on published datasets identifies ncRNA tails and compresses them into a human readable, portable, CSV format

To develop and validate this workflow, we used global 3’ end sequencing data from two previously published datasets, Labno et al. 2016 (hereafter called the Labno dataset) which investigated targets of human DIS3L2, and Son et al. 2018 (the Son dataset) which investigated targets of human PARN and TOE1. We also used a gene-specific 3’ end sequencing dataset from Lardelli and Lykke-Andersen 2020 (the Lardelli dataset), which investigated snRNA targets of human TOE1. After producing SAM alignment files using STAR with settings discussed above, Tailer, using a full genome annotation file (Ensembl 104), reduced gigabyte (Gb)-sized SAM files into megabyte (Mb)-sized Tail CSV files (Table 1), which makes uploading and analyzing on a web server practical. Output Tail CSV file sizes can be further reduced by using subset annotations with only genes of interest. The Tail CSV file can be used directly for visualization or analysis for users experienced with this type of data, or it can be fed into the Tailer-Analysis web app described below, or used directly in R with individual functions available from the GitHub repository.

**Table 1.**
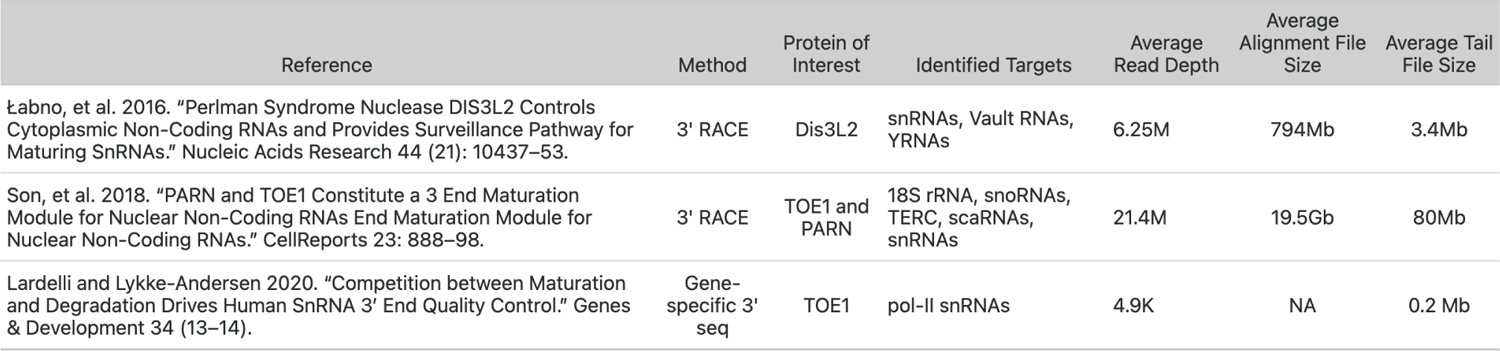
Dataset Summary

| Table 1: Dataset Summary |  |  |  |  |  |  |
| --- | --- | --- | --- | --- | --- | --- |
| Reference | Method | Protein of Interest | Identified Targets | Average Read Depth | Average Alignment File Size | Average Tail File Size |
| Labno, et al. 2016. "Perlman Syndrome Nuclease DIS3L2 Controls Cytoplasmic Non-Coding RNAs and Provides Surveillance Pathway for Maturing SnRNAs." Nucleic Acids Research 44 (21): 10437–53. | 3' RACE | Dis3L2 | snRNAs, Vault RNAs, YRNAs | 6.25M | 794Mb | 3.4Mb |
| Son, et al. 2018. "PARN and TOE1 Constitute a 3 End Maturation Module for Nuclear Non-Coding RNAs End Maturation Module for Nuclear Non-Coding RNAs." CellReports 23: 888–98. | 3' RACE | TOE1 and PARN | 18S rRNA, snoRNAs, TERC, scaRNAs, snRNAs | 21.4M | 19.5Gb | 80Mb |
| Lardelli and Lykke-Andersen 2020. "Competition between Maturation and Degradation Drives Human SnRNA 3' End Quality Control." Genes & Development 34 (13–14). | Gene-specific 3' seq | TOE1 | pol-II snRNAs | 4.9K | NA | 0.2 Mb |

### Tailer-Analysis: A Shiny webapp for candidate discovery and 3’ end data visualization

Tailer-Analysis provides a simple, user-friendly, and open-source GUI and is built using the R Shiny library (Chang et al. 2021). As an input, this analysis app takes the Tail CSV files generated from Tailer-Processing as described above. Multiple tail files can be uploaded in the “Tail File Upload” tab, including different experimental conditions to be compared, and experimental replicates (Fig. 3A). Using the table interface, users can enter metadata information which will group and average replicates, allowing for easy downstream comparisons and statistical analyses.

**FIGURE 3.**
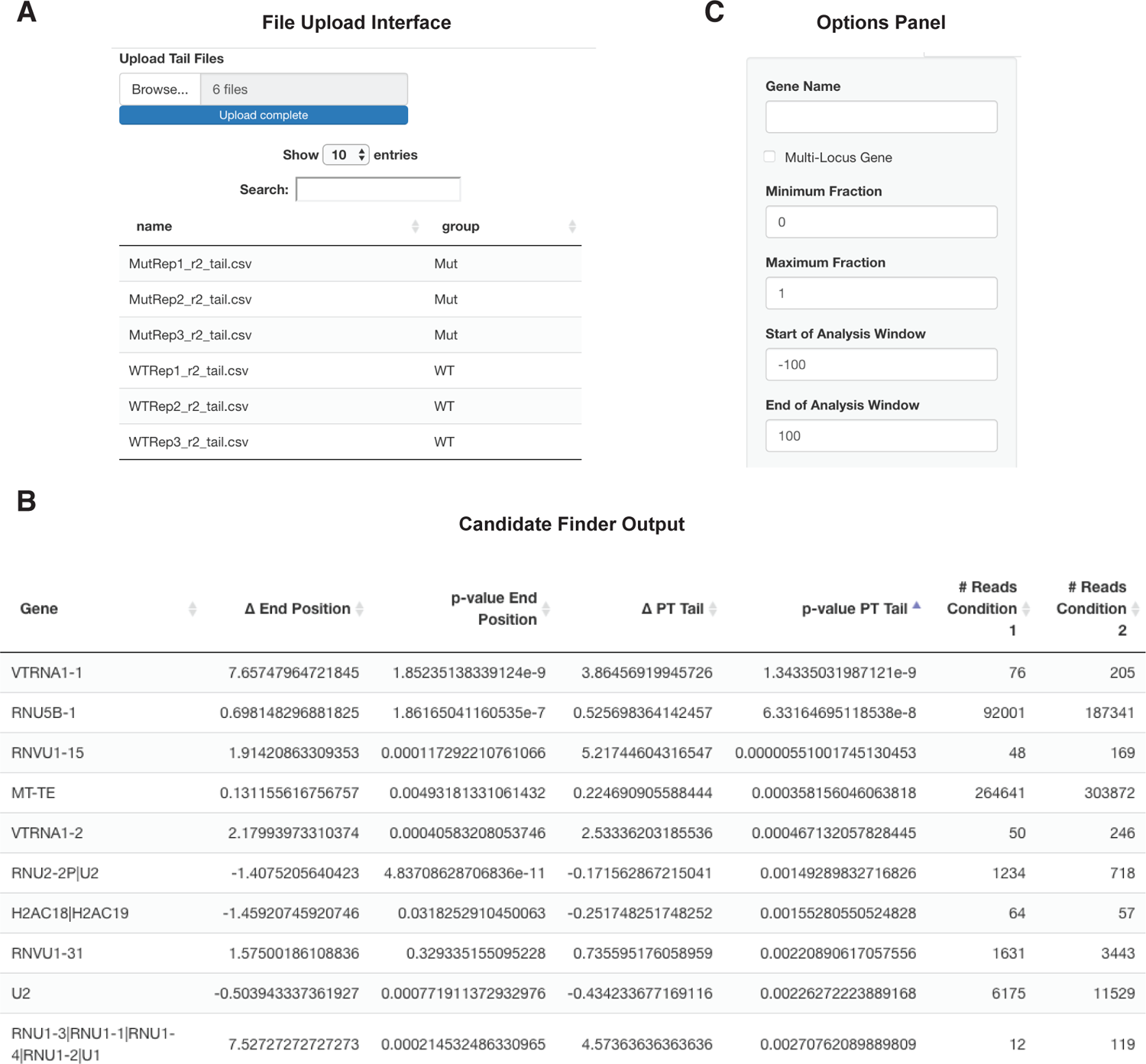
Example screenshots of the Tailer-Analysis Shiny interface. (A) Individual sample tail files can be uploaded to the webserver. Using the table interface, users can set grouping metadata, which will be used to bin replicates. After selecting the format data option, the user is provided with feedback on the conditions provided and number of samples. The user is also able to alter the order in which samples will be displayed using a simple drag and drop interface. (B) After uploading and setting metadata, users can use the Candidate Finder tab to rank RNAs based on their changes in tailing between the uploaded datasets. Reads can be filtered by minimum number of observations, magnitude of difference between conditions, and p-value. Hits are reported and ordered initially by statistical changes in RNA 3’ end positions but can also be re-ordered by statistical changes in post-transcriptional tail length by selecting the corresponding column. The candidate data can be downloaded and saved as a CSV file. (C) Every graph page contains an options side panel, which can be used to set the desired gene to be graphed and set different parameters for graphing. A checkbox for multi-locus genes is available to enable a slower but more accurate analysis of RNAs produced from multiple loci (see also Fig. 6 below).

On the “Candidate Finder” tab, users can compare two of their experimental conditions (Fig. 3B). RNAs with significant changes at their 3’ ends are found by comparing two conditions to look for statistically significant differences. The app first generates a list of all genes identified by Tailer-Processing. For each gene, the data from each replicate is then pooled and 3’ end positions and tail lengths are compared between experimental conditions using a Kolmogorov-Smirnov (KS) test. Candidate genes are reported in order of p-value for changes in 3’ end position but can also be sorted based on changes in tail length. This helps distinguish between conditions that may be affecting the trimming or extension of RNA molecules in general, versus conditions that affect post-transcriptionally added nucleotides specifically. Candidates can be filtered by minimum number of observations, p-value, and magnitude of difference in tail lengths.

Tail files generated from the Labno dataset were uploaded to the Shiny App web server and binned into a WT or a Mutant DIS3L2 condition. Using the built-in candidate discovery tool, a list of candidates with a minimum of 10 observed reads was generated (Supplemental Table 1). Among the top candidates with significantly altered 3’ ends were Vault RNA-1 (VTRNA-1), Y3 RNA (RNY3), and U6atac snRNA (U6ATAC), all of which were identified by Labno et al. Similarly, tail files from the Son dataset were subjected to candidate analysis using the Tailer-Analysis webapp. Identified potential targets (Supplemental Table 2) included many snoRNAs and scaRNAs which were targets also identified by Son et al. Thus, the Tailer pipeline faithfully recapitulates the identification of small RNA targets of 3’ end processing enzymes from published studies.

### Rapid visualization of 3’ end dynamics with the Tailer-Analysis webapp

The remaining tabs in the shiny app each correspond to graphs that can be used to individually explore RNA 3’ ends. 3’ ends of individual RNAs, either identified from the candidate discovery tool or of specific interest to the user, can be visually analyzed and compared between experiments as described in more detail below. These graph functions are written in ggplot2 (Wickham 2016; R Core Team 2021). For each graph, position 0 corresponds to the annotated mature RNA 3’ end. In cases where the mature 3’ ends of RNAs are incorrectly annotated in the provided annotation file, the position of the mature 3’ end can be manually adjusted using the options panel (Fig. 3C). Each plot also has an analysis window option whereby the user can limit their analysis to specific windows of 3’ end mapping. This can be used to exclude potential truncated RNAs from the analysis. Plotting and examining individual RNAs of interest can help distinguish between spurious hits and actual biological targets and can confirm that length changes are in the predicted direction and are of a sufficient magnitude to warrant further investigation.

The first two graphs visualize 3’ end positions of the sequenced population of the selected RNA. A bar graph gives a distribution of where the 3’ ends of the sequence reads are mapping in relation to the annotated mature 3’ end (Fig. 4A-C). Grey bars represent the positions, as fractions of the overall population, of the last genome-encoded nucleotide of the plotted RNA population. The colored bars represent the fraction of RNA molecules that contain post-transcriptionally added nucleotides at the indicated positions, broken down by nucleotide identity. It is important to emphasize, as detailed above, that post-transcriptional tails predicted by Tailer are the most conservative post-transcriptional tails based on the alignments.

**FIGURE 4.**
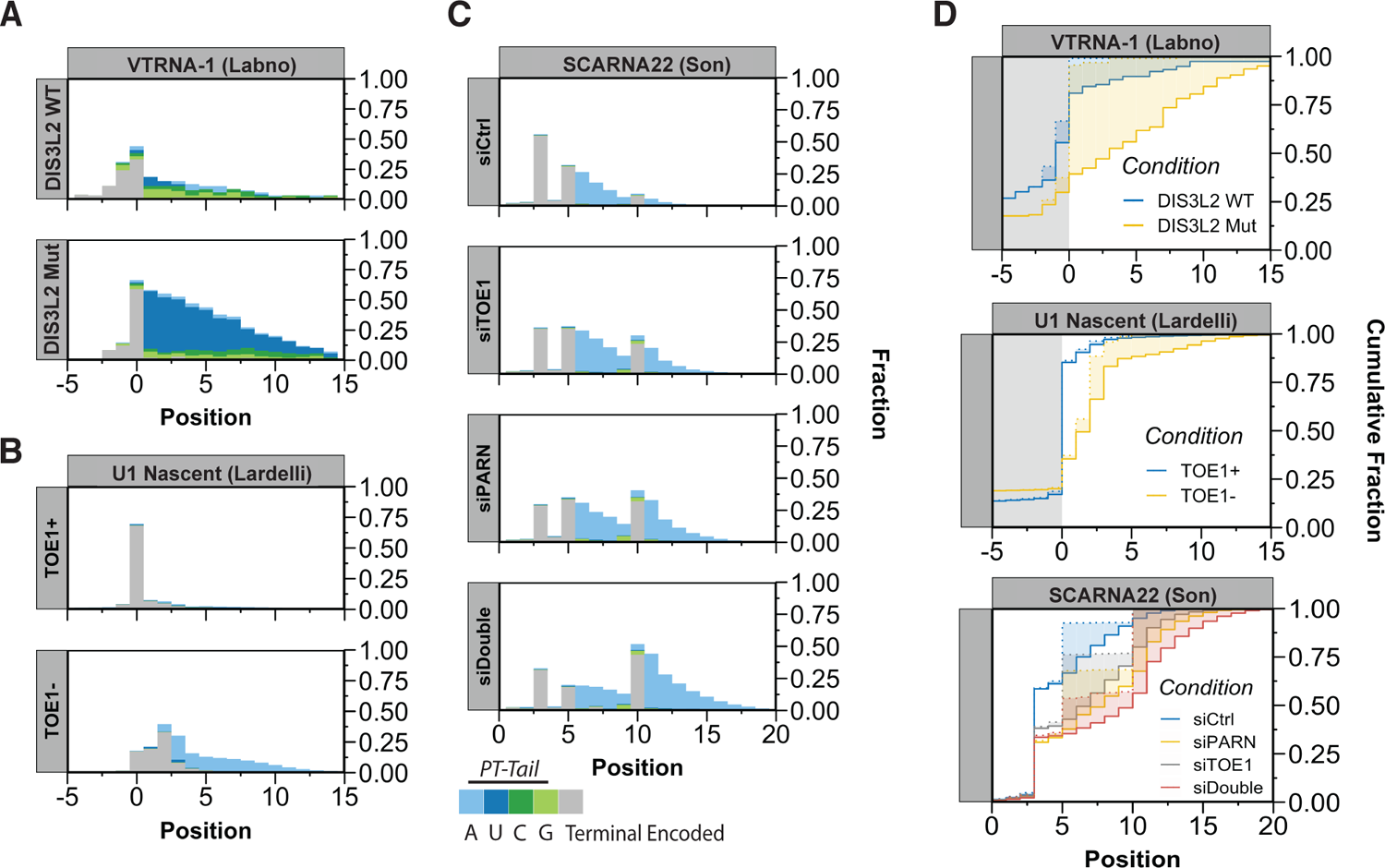
Sample plots of RNA 3’ end dynamics in response to processing factor depletion. (A-C) Tail bar graphs for indicated RNAs from the Labno (*A*), Lardelli (*B*) and Son (*C*) datasets, with grey bars showing the position of the terminal genomic encoded nucleotide as a fraction of the RNA population, and stacked colored bars showing the fraction of the RNA population containing post-transcriptional nucleotides at the indicated positions. (D) Cumulative plots displaying the cumulative fraction of overall 3’ end positions of the indicated RNA populations (including any post-transcriptional tail; solid lines) and the 3’ terminal genome-encoded nucleotide (dotted lines) with shading in between indicating the extent of post-transcriptional tailing. Dots to mark individual experiments can be toggled on and off using the options panel (Supplemental Figure 1).

Plotting VTRNA-1 from the Labno dataset recapitulates the presence of a post-transcriptional tail that consists primarily of uridines (dark blue bars) in the absence of DIS3L2 activity, which is observed on about half of the population and extends from the mature 3’ end (position 0), ranging from one to over ten uridines (Fig. 4A). Plotting U1 snRNA from the Lardelli dataset demonstrates accumulation of extended U1 snRNAs that are partially tailed with adenosines (light blue bars) in the absence of TOE1 (Fig. 4B). Furthermore, plotting SCARNA-22 from the Son data recapitulates the accumulation of extended RNA species terminating at the +10 position that accumulate with oligo-adenosine tails upon PARN and TOE1 depletion and a synergistic extension when both are depleted (Fig. 4C).

The second graph is a cumulative plot, which shows the cumulative fraction of RNA reads that map to specific 3’ end positions (Fig. 4D). Solid lines represent cumulative 3’ end positions of the RNA population when including post-transcriptional tails, and dotted lines represent the predicted 3’ ends excluding the post-transcriptional tails, with shading in between representing the extent of post-transcriptional tailing. The cumulative plots are particularly useful for comparing effects of different experimental conditions on specific RNAs in single graphs. Visualizing VTRNA1, SCARNA22 and U1 snRNA using this tool in Figure 4D recapitulates the overall extension of these transcripts upon depletion of the respective exonucleases, as observed by the overall right-shifts of the corresponding step plots. It can also be seen, by examining the extent of shading, that in the case of DIS3L2 inactivation (top panel) most of the difference in the VTRNA-1 3’ end is accounted for by differences in post-transcriptional tailing, consistent with the observations from the original study, whereas for PARN and TOE1 depletion (bottom two panels), effects are seen on both genome-encoded and post-transcriptional nucleotides of the target RNAs consistent with these enzymes trimming both post-transcriptional tails and genome-encoded nucleotides (Lardelli and Lykke-Andersen 2020; Son et al. 2018).

### Using Tailer-Analysis to visualize post-transcriptional tails

The next set of graphs focus on information concerning predicted post-transcriptionally added tails. The first graph is a logo plot containing information about proportions and compositions of post-transcriptional tails, with the 1 position corresponding to the first nucleotide of the tail and the height of each nucleotide representing the fraction of the RNA population that contains the modification (Fig. 5A-C). Plotting the datasets from Figure 4 in this manner reveals oligo-U tails that accumulate on VTRNA-1 in the absence of DIS3L2 activity (Fig. 5A) and oligo-A tails that accumulate on U1 snRNA (Fig. 5B) and SCARNA22 (Fig. 5C) in the absence of TOE1 and/or PARN activities. A background of primarily guanosines (denoted by a star in Fig. 5A) observed on VTRNA-1 appears, upon inspection of individual reads, to originate from an unknown linker in the Labno dataset.

**FIGURE 5.**
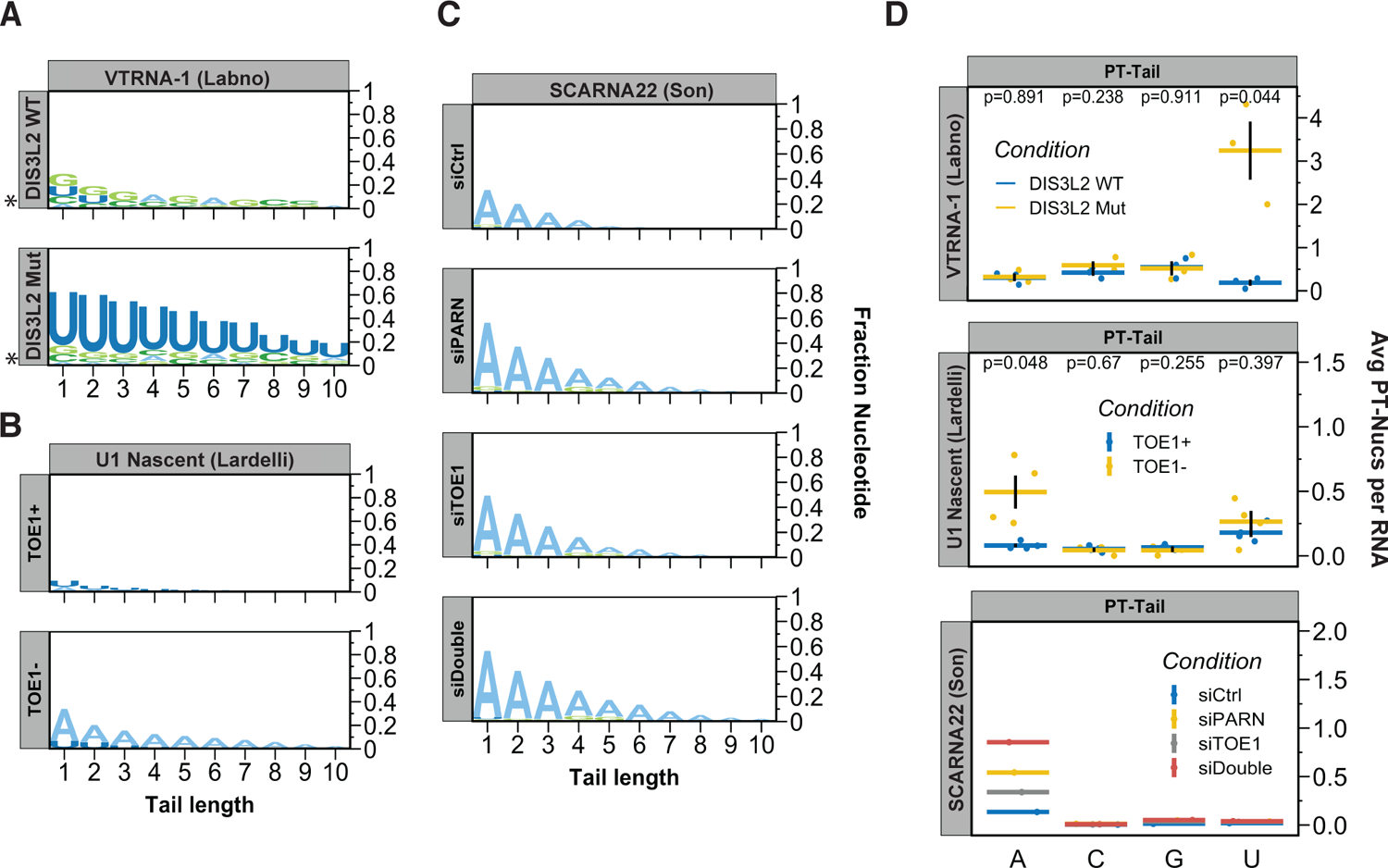
Sample plots of post-transcriptional tailing created by Tailer-Analysis. (A-C) Logo plots showing compositions of indicated RNA post-transcriptional tails as a fraction of the overall population. (D) Graphs showing average number of individual post-transcriptional nucleotides per RNA molecule as horizontal bars, with values for individual experiments shown by dots. In cases where multiple replicates are included (i.e. the Labno and Lardelli datasets), vertical lines show standard error of the mean (SEM) between experiments, and p-values from Student’s T-tests are reported to monitor significance.

The final graph shows the average number of post-transcriptional adenosines, uridines, guanosines, and cytidines found per read for the RNA of interest (Fig. 5D). In cases where replicates are included, this graph will show dots representing each experiment, bars for standard deviation, and, optionally, a p-value from a Student’s T-test. When applied to the analyzed datasets, these graphs again highlight the U-tailing observed for VTRNA-1, and A-tailing for SCARNA22 and U1 snRNAs upon depletion of the respective exonucleases. The source R code for generating all four types of graphs are available from our GitHub repository, are well documented, and can be imported and used in an active R session. Furthermore, below each graph is an option to download the raw plot data in CSV format. This option facilitates graphing using alternative software.

### Statistical outputs and outputs of values for manual graphing

The final tab of Tailer-Analysis contains utilities for testing statistical significance between groups (Supplementary Figure 2). After selecting two conditions to compare, the user is presented with tables of pair-wise KS-tests between each replicate in each condition. Statistical testing is done for both overall end position and total post-transcriptional tail length, which, as noted above, can help to distinguish between perturbations that affect post-transcriptional tailing only and those that also affect genome encoded tails. The page will also output a KS-test after pooling all replicates in each condition.

### Inclusive alignments to multi-loci genes prevents spurious tailing calls

A subset of small ncRNAs are produced from multiple loci in the genome, which in many cases are identical to one another except for their downstream sequences. Forcing a multi-locus RNA read to map to a single locus can lead to tails being falsely called that originate from the downstream sequence of a different locus from which the RNA was actually transcribed. As an example, human U1 snRNA originates from multiple active genes. Using any single locus in the Tailer analysis leads to the calling of spurious C- and U-post-transcriptional tails, which actually originate from other transcribed loci (Fig. 6A and B). In order to accurately assign reads, both global and local modes of Tailer allow for all loci to be considered when analyzing 3’ end tails, in which case the spuriously called C- and U-tailing of U1 snRNA is much reduced as reads are mapped to their proper loci (Fig. 6A). This demonstrates the importance of considering information from all gene loci when analyzing 3’ tailing data. Since post-transcriptional tail calls are conservative based on best alignment fits, analyses using multiple loci are more likely to miss a subset of actual short post-transcriptional tails, but, importantly, they help reduce the rate of false positive calls as observed for U1 snRNA.

**FIGURE 6.**
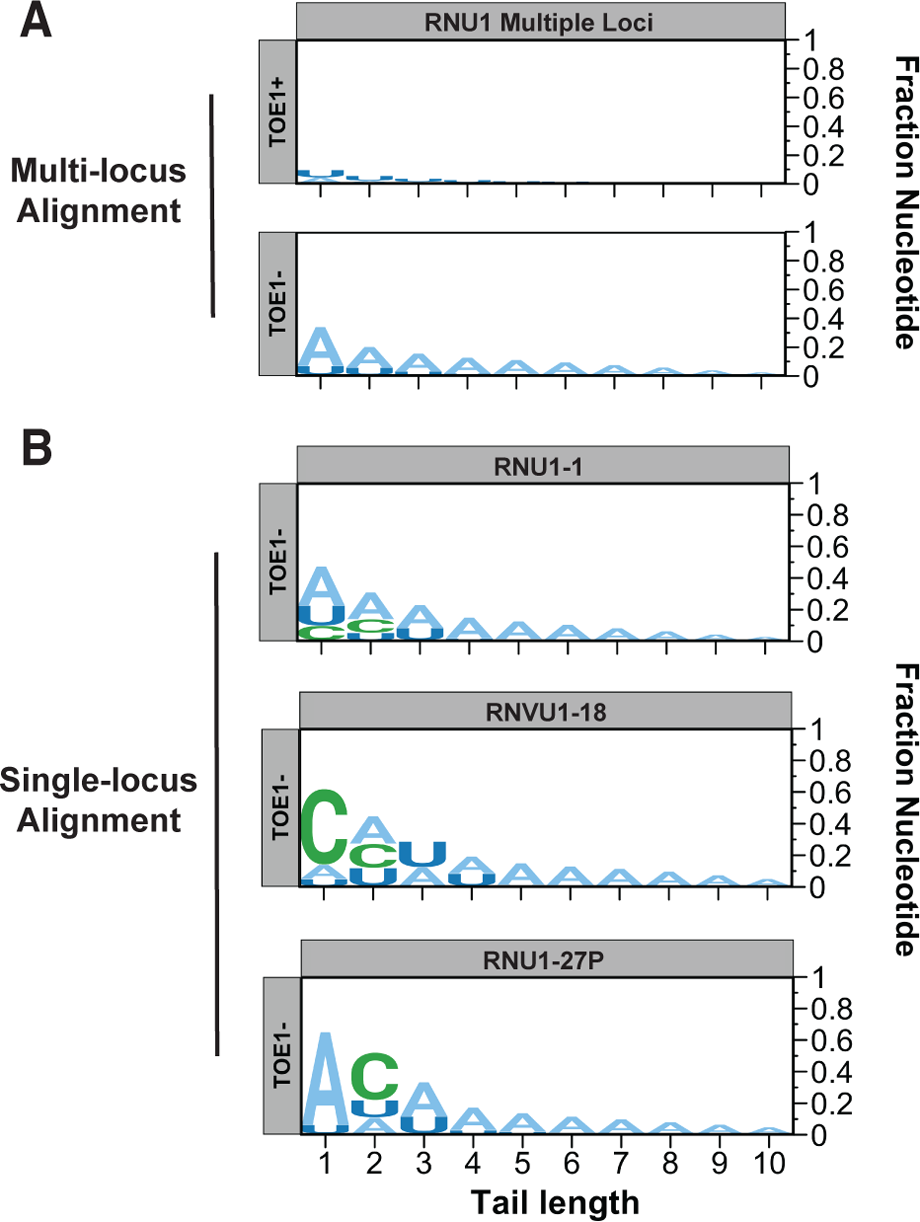
Logo plots of tails of a multi-locus RNA called with incomplete locus information. (A) U1 snRNAs from the Lardelli dataset aligned to all regular U1 loci of the genome, showing accumulation of post-transcriptional A-tails in the absence of TOE1. (B) The same dataset as in panel *A*, aligned to single U1 snRNA gene loci, leading to erroneous post-transcriptional tail calls.

## Conclusions

Trimming and tailing of RNA 3’ ends play key roles in the processing and quality control of non-coding RNAs. Advances in deep sequencing technologies and methods for library preparation have provided the tools to generate hordes of RNA 3’ end data. However, tools to analyze these types of data have remained limited. We developed Tailer to help spur inquiry into this important regulatory mechanism with a particular focus on ease of installation and use. Distribution of Tailer-Processing through PyPi allows for quick and easy installation in a wide variety of environments without end-users needing to manage dependencies and compatibility. Furthermore, users are not required to work with and manipulate genome or annotation data, such as making artificial genomes with singular loci for each gene. Tailer-Analysis as a web server allows users with no experience in R or coding to upload and explore datasets, and open-source distribution of the code allows for more advanced users to work more rapidly with their data using R.

It is important to note that with current protocols, because RNA 3’ end information is typically preserved using a ligation step, biases in the data are likely introduced due to effects of RNA structure or sequence on ligation efficiencies (Fuchs et al. 2015). Furthermore, certain RNAs terminate with a 3’ end modification that inhibits ligation (e.g. U6 snRNA terminating in a 2’-3’ cyclic phosphate) (Gu et al. 1997; Honda et al. 2016), which, if such modifications are not removed prior to ligation, may exclude that RNA entirely from analysis, or bias the analysis towards a state of maturation that does not contain the modification. In addition, since post-transcriptional tails are distinguished from nucleotides introduced during transcription by genome alignment, post-transcriptional tails can be missed, particularly short ones. Thus, analysis strategies based on ligation-mediated sequencing lend themselves best to monitoring changes to 3’ ends of RNAs between different cellular conditions (such as depletion of processing factors), rather than measuring accurate levels of tailing of one RNA over another. However, the development of direct RNA sequencing methods (Byrne et al. 2017), which can also readily be analyzed by the tools developed here, promise to alleviate many of these concerns.

Through a combined approach of local and global alignment, Tailer can, reproducibly and transparently, address many of the issues common with working with non-coding RNA sequencing data including the analysis of RNAs produced from multiple loci. The Tailer suite is validated by extensive analysis of datasets from three different studies using three different methods of library preparation. Thus, Tailer should allow more researchers to enter this research space and improve our understanding of this important mechanism of RNA regulation.

## Materials and Methods

### Tailer-Processing and Tailer-Analysis access

Source code for Tailer-Processing, the Tailer-Analysis Shiny app, further examples, and usage instructions can be found on our GitHub (https://github.com/TimNicholsonShaw/tailer and https://github.com/TimNicholsonShaw/tailer-analysis) and are available for use under the MIT license. Tailer-Analysis is available as a web server at https://timnicholsonshaw.shinyapps.io/tailer-analysis/.

### Data pre-processing

Data for these analyses was obtained from the NCBI GEO repository using the FASTQ-dump utility (Labno: GSE82336, Son: GSE111511, Lardelli: GSE141709). The Labno dataset was aligned without modification using the STAR aligner as described above. The Son dataset contained a four nucleotide 3’ adapter sequence which was trimmed using the FASTQ/A Trimmer from the FASTX-Toolkit and then aligned as above. The Lardelli dataset needed to have a 13-nucleotide barcode trimmed from the 3’ end which was performed using options provided by Tailer-Processing’s local mode and described in more detail in the repository’s readme (-x 13 flag).

### Tailer-Processing global

Global Tailer-Processing begins by generating a Search Query Language (SQL) database of all genes in the annotation file using the GFFutils module which allows for rapid look up. Tailer then reads in the provided SAM/BAM file using the PySam module, iterates through every read, discarding members of pairs that originate from the 5’ end of the RNA, and tags them with every gene that they overlap with in all their possible alignments using the SQL database, while collapsing identical reads together. For each aligned gene, tail position and composition are inferred using the soft-clipping flag in the CIGAR string of the SAM/BAM file and the annotated 3’ end of the gene from the GTF annotation file. Tail information is compared for all possible genes and the gene that gives an alignment closest to the annotated 3’ mature end is reported. In cases where multiple genes produce identical tail information, all genes are reported. The resulting tail information is then written to a CSV file referred to as a Tail CSV file.

### Tailer-Processing local

If provided with Ensembl IDs of interest, Tailer-Processing in local mode will contact Ensembl servers (requiring an internet connection), download their gene sequences via Ensembl’s REST API, and build a FASTA file. If provided with a reference FASTA file, it will instead use that reference, which should include downstream sequence for accurate distinction between encoded and post-transcriptional tails, in which case the length of the downstream sequence should be indicated with the -m flag for accurate annotation of the mature 3’ end. This is done automatically when providing an Ensembl ID. The mature end can also be adjusted later in the options panel of Tailer-Analysis. Using the command line BLAST utility makeBlastDb, Tailer creates a database compatible with BLAST searches. Tailer then uses the query FASTQ to generate a BLAST compatible query file and, using command-line blast, searches the query against the reference outputting the results in JSON format. After parsing the output, Tailer infers tails for each aligned read using alignment to the reference sequence and reports the tail for the gene(s) whose 3’ end is closest to the 3’ end of the gene. The resulting tail information is then written to a Tail CSV file.

### Tailer-Analysis

Tailer-Analysis takes Tail CSV files generated above and metadata provided by the user indicating replicate groups and creates a singular data frame in long format (1 observation per row). This data frame is then fed into the other tools provided by Tailer-Analysis. For the candidate finder, replicates from two different groups are pooled and compared with a K-S test which is reported for End Position and for Tail Length. This list is sorted by End Position P-value and reported to the user. Tail bar graphs are initiated by creating a matrix of frequencies of each nucleotide or genome encoded end at every requested position. This matrix is fed to ggplot’s geom_bar function and faceted based on the experimental condition. Cumulative plots are created by calculating cumulative sums at each position for both End Position (total tail length with post-transcriptional additions) and End Position minus Tail Length (location of the genome-encoded end).

This data is summarized and averaged based on condition and position and fed to a geom_step ggplot function. The tail logo grapher calculates nucleotide frequencies at all requested positions and feeds the frequency matrix to ggseqlogo’s geom_logo function (Wagih 2017). The post-transcriptional nucleotide graph is created by first finding the number of each nucleotide in the Tail Sequence column for each sample and calculating the mean count of each nucleotide per RNA molecule. Data is then summarized based on condition and nucleotide and fed to ggplot using geom_jitter for dots, geom_segment for lines, and geom_errorbar (SEM reported). Uniform theming is accomplished with a single defined common theme that is applied to all graphs and can be reviewed on the GitHub repository for Tailer-Analysis.

## Acknowledgements

We would like to thank TSCC at the San Diego Computing Center for use of their hardware for alignments. We would also like to thank Dr. Elly Poretsky for enlightening conversations concerning Shiny apps, Dr. Brian Tsu for Pythonic-based encouragement and general enthusiasm, and members of the Lykke-Andersen lab, Alberto Carreño, Cody Ocheltree, and Tiantai Ma for feedback and testing. This work was supported by National Institutes of Health (NIH) grant R35 GM118069 awarded to J. L.-A.

## References

1. Allmang C, Kufel J, Chanfreau G, Mitchell P, Petfalski E, Tollervey D. 1999. Functions of the exosome in rRNA, snoRNA and snRNA synthesis. EMBO J 18: 5399–410.

2. Berndt H, Harnisch C, Rammelt C, Stöhr N, Zirkel A, Dohm JC, Himmelbauer H, Tavanez JP, Hüttelmaier S, Wahle E. 2012. Maturation of mammalian H/ACA box snoRNAs: PAPD5-dependent adenylation and PARN-dependent trimming. RNA 18: 958–972.

3. Byrne A, Beaudin AE, Olsen HE, Jain M, Cole C, Palmer T, DuBois RM, Forsberg EC, Akeson M, Vollmers C. 2017. Nanopore long-read RNAseq reveals widespread transcriptional variation among the surface receptors of individual B cells. Nat Commun 8.

4. Chang W, Cheng J, Allaire JJ, Sievert C, Schloerke B, Xie Y, Allen J, McPherson J, Dipert A, Borges B. 2021. shiny: Web Application Framework for R. R package version 1.6.0. https://CRAN.R-project.org/package=shiny

5. Darnell JE, Philipson L, Wall R, Adesnik M. 1971. Polyadenylic acid sequences: Role in conversion of nuclear RNA into messenger RNA. Science 174: 507–510.

6. Deutscher MP. 1973. Synthesis and Functions of the -C-C-A Terminus of Transfer RNA. Prog Nucleic Acid Res Mol Biol 13: 51–92.

7. Dobin A, Davis CA, Schlesinger F, Drenkow J, Zaleski C, Jha S, Batut P, Chaisson M, Gingeras TR. 2013. STAR: ultrafast universal RNA-seq aligner. Bioinformatics 29: 15–21.

8. Dupasquier M, Kim S, Halkidis K, Gamper H, Hou YM. 2008. tRNA Integrity Is a Prerequisite for Rapid CCA Addition: Implication for Quality Control. J Mol Biol 379: 579–588.

9. Edmonds M, Vaughan MH, Nakazato H. 1971. Polyadenylic acid sequences in the heterogeneous nuclear RNA and rapidly-labeled polyribosomal RNA of HeLa cells: possible evidence for a precursor relationship. Proc Natl Acad Sci U S A 68: 1336– 1340.

10. Frohman MA, Dush MK, Martin GR. 1988. Rapid production of full-length cDNAs from rare transcripts: Amplification using a single gene-specific oligonucleotide primer. Proc Natl Acad Sci U S A 85: 8998–9002.

11. Fuchs RT, Sun Z, Zhuang F, Robb GB. 2015. Bias in Ligation-Based Small RNA Sequencing Library Construction Is Determined by Adaptor and RNA Structure. PLoS One 10: e0126049.

12. Goldstrohm AC, Wickens M. 2008. Multifunctional deadenylase complexes diversify mRNA control. Nat Rev Mol Cell Biol 9: 337–344.

13. Gu J, Shumyatsky G, Makan N, Reddy R. 1997. Formation of 2’,3’-cyclic phosphates at the 3’ end of human U6 small nuclear RNA in vitro: Identification of 2’,3’-cyclic phosphates at the 3’ ends of human signal recognition particle and mitochondrial RNA processing RNAs. J Biol Chem 272: 21989–21993.

14. Honda S, Morichika K, Kirino Y. 2016. Selective amplification and sequencing of cyclic phosphate–containing RNAs by the cP-RNA-seq method. Nat Protoc 11: 476–489.

15. Howe KL, Achuthan P, Allen J, Allen J, Alvarez-Jarreta J, Ridwan Amode M, Armean IM, Azov AG, Bennett R, Bhai J, et al. 2021. Ensembl 2021. Nucleic Acids Res 49: D884–D891.

16. Katoh T, Sakaguchi Y, Miyauchi K, Suzuki T, Suzuki T, Kashiwabara SI, Baba T. 2009. Selective stabilization of mammalian microRNAs by 3’ adenylation mediated by the cytoplasmic poly(A) polymerase GLD-2. Genes Dev 23: 433–438.

17. Łabno A, Warkocki Z, Kulínski T, Krawczyk PS, Bijata K, Tomecki R, Dziembowski A. 2016. Perlman syndrome nuclease DIS3L2 controls cytoplasmic non-coding RNAs and provides surveillance pathway for maturing snRNAs. Nucleic Acids Res 44: 10437–10453.

18. LaCava J, Houseley J, Saveanu C, Petfalski E, Thompson E, Jacquier A, Tollervey D. 2005. RNA Degradation by the Exosome Is Promoted by a Nuclear Polyadenylation Complex. Cell 121: 713–724.

19. Lardelli RM, Lykke-Andersen J. 2020. Competition between maturation and degradation drives human snRNA 3′ end quality control. Genes Dev 34: 1–13.

20. Lardelli RM, Schaffer AE, C Eggens VR, Zaki MS, Grainger S, Sathe S, Van Nostrand EL, Schlachetzki Z, Rosti B, Akizu N, et al. 2017. Biallelic mutations in the 3′ exonuclease TOE1 cause pontocerebellar hypoplasia and uncover a role in snRNA processing. Nat Genet 49: 457–466.

21. Lee M, Choi Y, Kim K, Jin H, Lim J, Nguyen TA, Yang J, Jeong M, Giraldez AJ, Yang H, et al. 2014. Adenylation of maternally inherited MicroRNAs by wispy. Mol Cell 56: 696–707.

22. Lee SY, Mendecki J, Brawerman G. 1971. A polynucleotide segment rich in adenylic acid in the rapidly-labeled polyribosomal RNA component of mouse sarcoma 180 ascites cells. Proc Natl Acad Sci U S A 68: 1331–1335.

23. Li H, Handsaker B, Wysoker A, Fennell T, Ruan J, Homer N, Marth G, Abecasis G, Durbin R. 2009. The Sequence Alignment/Map format and SAMtools. Bioinformatics 25: 2078–2079.

24. Liu X, Zheng Q, Vrettos N, Maragkakis M, Alexiou P, Gregory BD, Mourelatos Z. 2014. A MicroRNA Precursor Surveillance System in Quality Control of MicroRNA Synthesis. Mol Cell 55: 868–879.

25. Liudkovska V, Dziembowski A. 2021. Functions and mechanisms of RNA tailing by metazoan terminal nucleotidyltransferases. Wiley Interdiscip Rev RNA 12e1622

26. Lund E, Dahlberg JE. 1992. Cyclic 2′,3′-phosphates and nontemplated nucleotides at the 3′ end of spliceosomal U6 small nuclear RNA’s. Science 255: 327–330.

27. Nguyen D, Grenier St-Sauveur V, Bergeron D, Dupuis-Sandoval F, Scott MSS, Bachand F. 2015. A Polyadenylation-Dependent 3′ End Maturation Pathway Is Required for the Synthesis of the Human Telomerase RNA. Cell Rep 13: 2244– 2257.

28. Nicholson AL, Pasquinelli AE. 2019. Tales of Detailed Poly(A) Tails. Trends Cell Biol 29: 191–200.

29. Perumal K, Reddy R. 2002. The 3′ end formation in small RNAs. Gene Expr 10: 59–78.

30. Pirouz M, Ebrahimi AG, Gregory RI. 2019. Unraveling 3’-end RNA uridylation at nucleotide resolution. Methods 155: 10–19.

31. Quinlan AR, Hall IM. 2010. BEDTools: A flexible suite of utilities for comparing genomic features. Bioinformatics 26: 841–842.

32. R Core Team. 2021. R: A language and environment for statistical computing. R Foundation for Statistical Computing, Vienna, Austria. URL https://www.R-project.org/.

33. Rinke J, Steitz JA. 1982. Precursor molecules of both human 5S ribosomal RNA and transfer RNAs are bound by a cellular protein reactive with anti-La Lupus antibodies. Cell 29: 149–159.

34. Shcherbik N, Wang M, Lapik YR, Srivastava L, Pestov DG. 2010. Polyadenylation and degradation of incomplete RNA polymerase I transcripts in mammalian cells. EMBO Rep 11: 106–11.

35. Shukla S, Parker R. 2017. PARN Modulates Y RNA Stability and Its 3′-End Formation. Mol Cell Biol 37: e00264–17.

36. Son A, Park JE, Kim VN. 2018. PARN and TOE1 Constitute a 3′ End Maturation Module for Nuclear Non-coding RNAs. Cell Rep 23: 888–898.

37. Suzuki S, Yasuda T, Shiraishi Y, Miyano S, Nagasaki M. 2011. ClipCrop: a tool for detecting structural variations with single-base resolution using soft-clipping information. BMC Bioinformatics 12: S7.

38. Van Rossum G, Drake FL. 2019. Python 3 Reference Manual. CreateSpace, Scotts Valley, CA.

39. Wagih O. 2017. ggseqlogo: A ‘ggplot2’ Extension for Drawing Publication-Ready Sequence Logos. R package version 0.1. https://CRAN.R-project.org/package=ggseqlogo

40. Wickham H. 2016. ggplot2: Elegant Graphics for Data Analysis. Springer-Verlag New York.

41. Wolin SL, Maquat LE. 2019. Cellular RNA surveillance in health and disease. Science 366: 822–827.

42. Yu S, Kim VN. 2020. A tale of non-canonical tails: gene regulation by post-transcriptional RNA tailing. Nat Rev Mol Cell Biol 2020 *219* 21: 542–556.

