## Supplementary material for "Tailer: A Pipeline for Sequencing-Based Analysis of Non-Polyadenylated RNA 3’ End Processing": Figure S1

**Supplementary Figure 1.** Cumulative plots with replicate dots

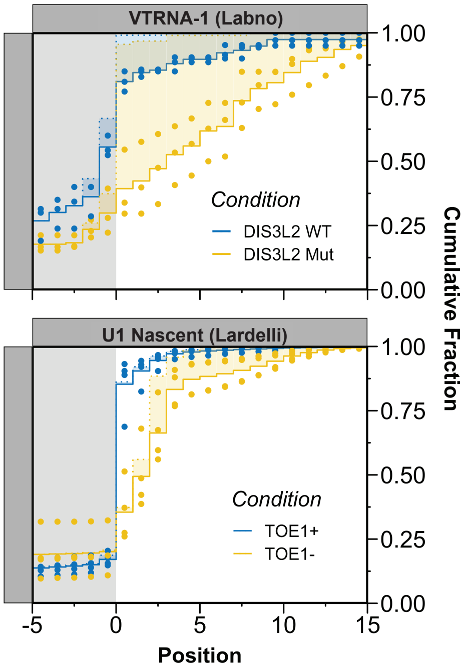


**Figure S1.** Recreation of cumulative plots with optional replicate dots. Replicate dots can be toggled on and off using the options panel.
