## Supplementary material for "Tailer: A Pipeline for Sequencing-Based Analysis of Non-Polyadenylated RNA 3’ End Processing": Figure S2

**Supplementary Figure 2.** Example statistics output

### VTRNA1-1 (Labno)

**End Position KS-test Matrix**

| | DIS3L2 WT_1 | DIS3L2 WT_2 | DIS3L2 WT_3 |
|---|---|---|---|
| DIS3L2 Mut_1 | 4.829023E-05 | 4.339632E-04 | 2.187131E-01 |
| DIS3L2 Mut_2 | 7.084303E-04 | 1.295540E-03 | 3.120647E-01 |
| DIS3L2 Mut_3 | 9.286356E-06 | 1.645482E-04 | 2.532410E-01 |

**Pooled End Position KS-test**

1.852351e-09

**Tail Length KS-test Matrix**

| | DIS3L2 WT_1 | DIS3L2 WT_2 | DIS3L2 WT_3 |
|---|---|---|---|
| DIS3L2 Mut_1 | 2.377555E-05 | 7.524774E-05 | 2.187131E-01 |
| DIS3L2 Mut_2 | 1.222320E-03 | 2.284022E-03 | 6.107870E-01 |
| DIS3L2 Mut_3 | 5.426585E-06 | 4.051894E-05 | 1.465849E-01 |

**Pooled Tail Length KS-test**

1.34335e-09

**Number of Observations**

| Condition | n |
|---|---|
| DIS3L2 WT_1 | 21 |
| DIS3L2 WT_2 | 20 |
| DIS3L2 WT_3 | 35 |
| DIS3L2 Mut_1 | 54 |
| DIS3L2 Mut_2 | 118 |
| DIS3L2 Mut_3 | 33 |

### RNU1-1 (Lardelli)

**End Position KS-test Matrix**

| | TOE1+_1 | TOE1+_2 | TOE1+_3 | TOE1+_4 |
|---|---|---|---|---|
| TOE1-_1 | <1e-50 | <1e-50 | <1e-50 | <1e-50 |
| TOE1-_2 | <1e-50 | <1e-50 | <1e-50 | <1e-50 |
| TOE1-_3 | <1e-50 | <1e-50 | 2.81840850657744e-09 | <1e-50 |
| TOE1-_4 | <1e-50 | <1e-50 | <1e-50 | <1e-50 |

**Pooled End Position KS-test**

<1e-50

**Tail Length KS-test Matrix**

| | TOE1+_1 | TOE1+_2 | TOE1+_3 | TOE1+_4 |
|---|---|---|---|---|
| TOE1-_1 | <1e-50 | <1e-50 | 3.30846461338297e-14 | 8.8159308742064e-09 |
| TOE1-_2 | <1e-50 | <1e-50 | 9.64485158405637e-11 | 4.19450031485802e-07 |
| TOE1-_3 | <1e-50 | <1e-50 | 0.000124527617351933 | 0.000211237539438636 |
| TOE1-_4 | <1e-50 | <1e-50 | 6.50890452646991e-12 | 2.96518487430397e-07 |

**Pooled Tail Length KS-test**

<1e-50

**Number of Observations**

| Condition | n |
|---|---|
| TOE1+_1 | 3495 |
| TOE1+_2 | 4576 |
| TOE1+_3 | 2567 |
| TOE1+_4 | 8044 |
| TOE1-_1 | 1704 |
| TOE1-_2 | 1355 |
| TOE1-_3 | 1644 |
| TOE1-_4 | 1285 |

**Figure S2.** Example output from the Tailer-Analysis' statistics tab. After selecting a gene of interest, each replicate in a condition is compared pairwise to every other replicate in the comparison condition using a KS test. Comparisons are performed for the End Position metric and the Tail Length metric described above. Statistics are also performed after pooling all samples to get a singular pooled KS test metric. The number of observations in each condition is also reported in a table.
